# Comparison of retinol binding protein 1 with cone specific G-protein as putative effector molecules in cryptochrome signalling

**DOI:** 10.1101/2024.08.27.609845

**Authors:** Chad Yee, Rabea Bartölke, Katharina Görtemaker, Jessica Schmidt, Bo Leberecht, Henrik Mouritsen, Karl-Wilhelm Koch

**Author notes:** to whom correspondence should be addressed: Department of Neuroscience, Division of Biochemistry, Carl von Ossietzky Universität Oldenburg, 26111 Oldenburg, Germany, Neurosensorics/Animal Navigation, Institute of Biology and Environmental Sciences, Carl von Ossietzky Universität Oldenburg D-26111 Oldenburg, Germany. These authors contributed equally to the work (Chad Yee, Rabea Bartölke).

## Abstract

Vision and magnetoreception in navigating songbirds are strongly connected as recent findings link a light dependent radical-pair mechanism in cryptochrome proteins to signalling pathways in cone photoreceptor cells. A previous yeast-two-hybrid screening approach identified six putative candidate proteins showing binding to cryptochrome type 4a. So far, only the interaction of the cone specific G-protein transducin α-subunit was investigated in more detail. In the present study, we compare the binding features of the G-protein α-subunit with those of another candidate from the yeast-two-hybrid screen, cellular retinol binding protein. Purified recombinant European robin retinol binding protein bound retinol with high affinity, displaying an EC_50_ of less than 5 nM, thereby demonstrating its functional state. We applied surface plasmon resonance and a Förster resonance transfer analysis to test for interactions between retinol binding protein and cryptochrome 4a. In the absence of retinol, we observed no robust binding events, which contrasts the strong interaction we observed between cryptochrome 4a and the G-protein α-subunit. We conclude that retinol binding protein is unlikely to be involved in the primary magnetosensory signalling cascade.

## Introduction

Growing interest in the field of biological magnetoreception has intensified research to uncover the cellular and molecular mechanism by which animals can sense and use the Earth’s magnetic field to navigate. A light-dependent radical-pair mechanism^1-3^ provides a model hypothesis to understand static and radiofrequency magnetic field effects observed in several species, in particular in night-migratory songbirds^4-10^. One of the promising leads thus far suggests that cryptochrome (Cry) proteins may be the primary magnetic sensor molecules in several species, as suggested by studies involving birds^11-17^, various insects^18-20^, but see ref.^21^, and fish^22,23^, which suggests that it possibly evolved on more than one occasion. Detailed *in-vitro* studies on purified cryptochromes showed that European robin Cry4a exhibits the photochemistry and magnetic sensitivity consistent with the role of a protein candidate capable of sensing magnetic fields^14,24,25^. Furthermore, immunohistochemical staining techniques localized Cry4 in long wavelength single and double cone outer segments from European robin^12,16^ and, recently, behavioral studies supported that the radical-pair mechanism is based on a flavin-bound magnetoreceptor protein^9,10^, which in turn provided more support for a Cry protein being involved in magnetoreceptive signaling in night-migratory songbirds. A deeper account of current discussions of this topic is provided in several reviews^3,26-29^.

If Cry4 is the magnetic sensor in many animals, then it must be able to generate a signal that reaches the nervous system. For this, it likely needs the assistance of some form of signaling cascade involving additional proteins that can interpret and amplify the signal. In the search for potential protein candidates that could fulfill this role, a yeast-two-hybrid screen against a European robin retina cDNA library was performed^30^. This study narrowed down the list of putative candidates to six proteins that appeared to bind Cry4 in a cellular system. The six candidates were grouped into three categories^30^. One includes proteins directly involved in the phototransduction pathway. A follow-up study confirmed the interaction of Cry4a with the cone-specific G-protein transducin alpha-subunit using purified proteins and different interaction analyses^31^. Other candidates from the study of Wu et al. (2020)^30^ were the cellular retinol binding protein 1 (ErRBP1) and the retinal G protein-coupled receptor RGR, which are involved in retinoid metabolism. The involvement of phototransduction specific proteins in light-dependent magnetoreception somehow seems reasonable, although a concrete mechanism is missing so far. However, the thought of a retinol binding protein playing a potential role in magnetic signaling is both intriguing and puzzling, and we therefore decided to investigate this potential interaction partner further.

Retinol binding proteins are best known for transporting retinol and retinoids, or vitamin A, which would otherwise suffer from limited solubility and stability within an aqueous environment^32^. In the retinol-RBP complex, the protein structure almost completely encapsulates the retinol^32,33^, allowing a safe delivery to and within cells, where it plays various critical roles in signaling, vision, development, and more^33-36^. To carefully regulate the availability of vitamin A, multiple retinol binding proteins have developed in humans, including a plasma specific variant, as well as at least five intracellular variants that bind the various retinoids with different affinities, all of which appear to be necessary for proper vitamin A metabolism^37^. Each variant appears to be quite important, as highly conserved homologs for each exist throughout the animal kingdom. With at least several variants likely present in the avian retina, it is interesting to note that only one single variant (ErRBP1) appeared in the list of potential ErCry4 interaction candidates in the study of Wu et al. (2020)^30^.

As stated above, at first glance, it is puzzling to think that a retinol binding protein could take up a role in magnetic signaling. Retinol binding proteins belong to the superfamily of lipocalin proteins, a nearly ubiquitous family known for binding and transporting hydrophobic substances within a beta-barrel structure^32^. However, they are also known for moonlighting, which includes performing enzymatic activities and interactions with other proteins, in addition to their traditional ligand-binding role. Even human retinol binding proteins themselves appear to take part in various protein-protein interactions^38,39^. This certainly allowed for some interesting speculation on how a retinol binding protein might have picked up a moonlighting role in magnetoreception, and the potential implications thereof.

In the present work, we performed a brief bioinformatic analysis of the protein sequence, expressed and purified the protein, and characterized its biological function. The biologically active ErRBP1 was then used in interaction studies with ErCry4a, for which the binding of the cone-specific G-protein alpha-subunit to Cry4a served as a positive control and benchmark.

## Results

### Bioinformatics analysis

Wu et al. (2020) identified the gene RBP1 coding for retinol binding protein 1 as a putative interaction partner of Cry4a^30^. We performed a sequence alignment of cellular retinol binding protein 1 from different species and clearly classified ErRBP1 as a homolog of cellular retinol binding protein 1 (CRBP1), possessing 84.33% sequence identity with human CRBP1, but only 54-55% sequence identity with other human cellular RBPs. When compared to the chicken (Gallus gallus) CRBP1, the sequence identity was over 95%, and for many newly annotated avian species it is even higher^40^. Using the full construct amino acid sequence (including a 6x His tag), a theoretical molar extinction coefficient of 26,470 M^-1^ cm^-1^ was suggested by the online ProtParam tool by Expasy (see Methods section), along with a theoretical molecular mass of 17,051 Dalton.

### Functional characterization of ErRBP1

For protein-protein interaction analyses using SPR spectroscopy it was necessary to express, purify and characterize the hypothetical binding partners ErRBP1 and ErCry4a. A full characterization of purified ErCry4a has been described recently^14^ and no new characterization was needed since we used ErCry4a from the same source in the present study. For ErRBP1 we performed a thorough analysis of its spectroscopic and retinol binding characteristics as outlined below.

The purified protein was free of contaminating bands in an SDS polyacrylamide gel (see Figure S1 in the supplement). A lack of absorbance above 300 nm furthermore suggests the absence of any retinoids or other light absorbing contaminants in our preparations (see bold black line in Figure S2). Correctly folded and biologically active ErRBP1 should bind retinol in a concentration dependent manner. Free retinol, when titrated into either buffer or 100% ethanol, has a peak absorbance at approximately 325 nm (red spectra in Figure 1). The absorbance of our ErRBP1 + retinol samples show a peak absorbance shifted to approximately 350 nm, which is expected for ErRBP1-bound retinol (black spectra in Figure 1) based on previous work on CRBP from liver^41^. Two shoulder peaks and an increase in the magnitude in the absorbance spectrum of retinol also become prominent upon binding to ErRBP1 in agreement with previous findings^41^. We extended the titration going beyond the apparent saturation around 11 µM retinol (Figure S2) and applied the titration method for determination of the molar extinction coefficients of ErRBP1 (ε_280_ = 25560 ± 1037 M^-1^cm^-1^), free retinol (ε_325_ = 26,100 ± 1045 M^-1^cm^-1^), and retinol bound to ErBP1 (ε_350_ = 50,960 ± 2690 M^-1^cm^-1^). A detailed description of the titration method is provided in the supplement.

**Figure 1.**
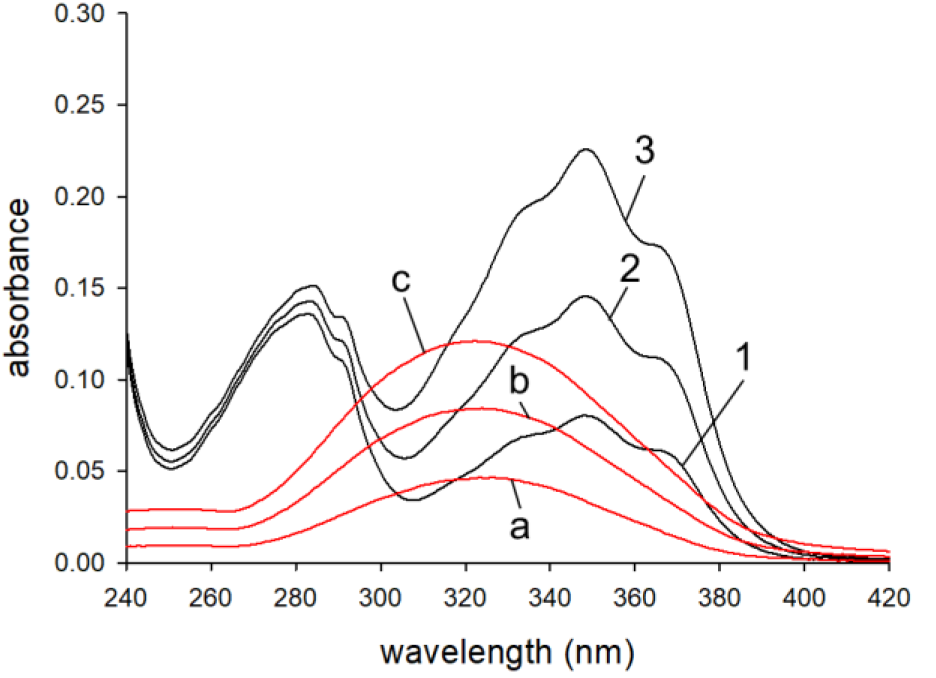
UV-vis Absorbance spectrum of purified ErRBP1 after consecutive additions of all-trans retinol. The protein concentration was ∼5.1 µM, and curves are shown for ∼1.57, 3.16, 4.73 µM added retinol (solid black lines corresponding to spectra 1, 2 and 3). The titration was repeated, but injecting retinol into buffer with identical concentrations (red lines corresponding to spectra a, b and c).

### Binding affinity of retinol to ErRBP1

A second crucial parameter for testing the biological function of ErRBP1 is the quantitative determination of ErRBP1’s binding affinity for retinol. Based on previous observations that fluorescence resonance energy transfer occurs from an excited tryptophan in RBP to bound retinol^42,43^, we set up an experiment using 500 nM ErRBP1 and titration with increasing concentrations of retinol. As expected, tryptophan fluorescence emission observed at 340 nm (upper spectrum in Figure 2A) decreased with increasing retinol concentrations (Figure 2A). Fluorescence emission of retinol was observed at 485 nm (see slight increase above 450 nm in Figure 2A). Several similar sets of data indicated that the retinol fluorescence intensity reached approximately 1/20^th^ of the intensity of the initial tryptophan fluorescence, and that the fluorescence intensity of retinol greatly increased upon binding to ErRBP1 from an aqueous environment, similar to previously seen results^42^. These results showed that purified ErRBP1 binds retinol.

**Figure 2.**
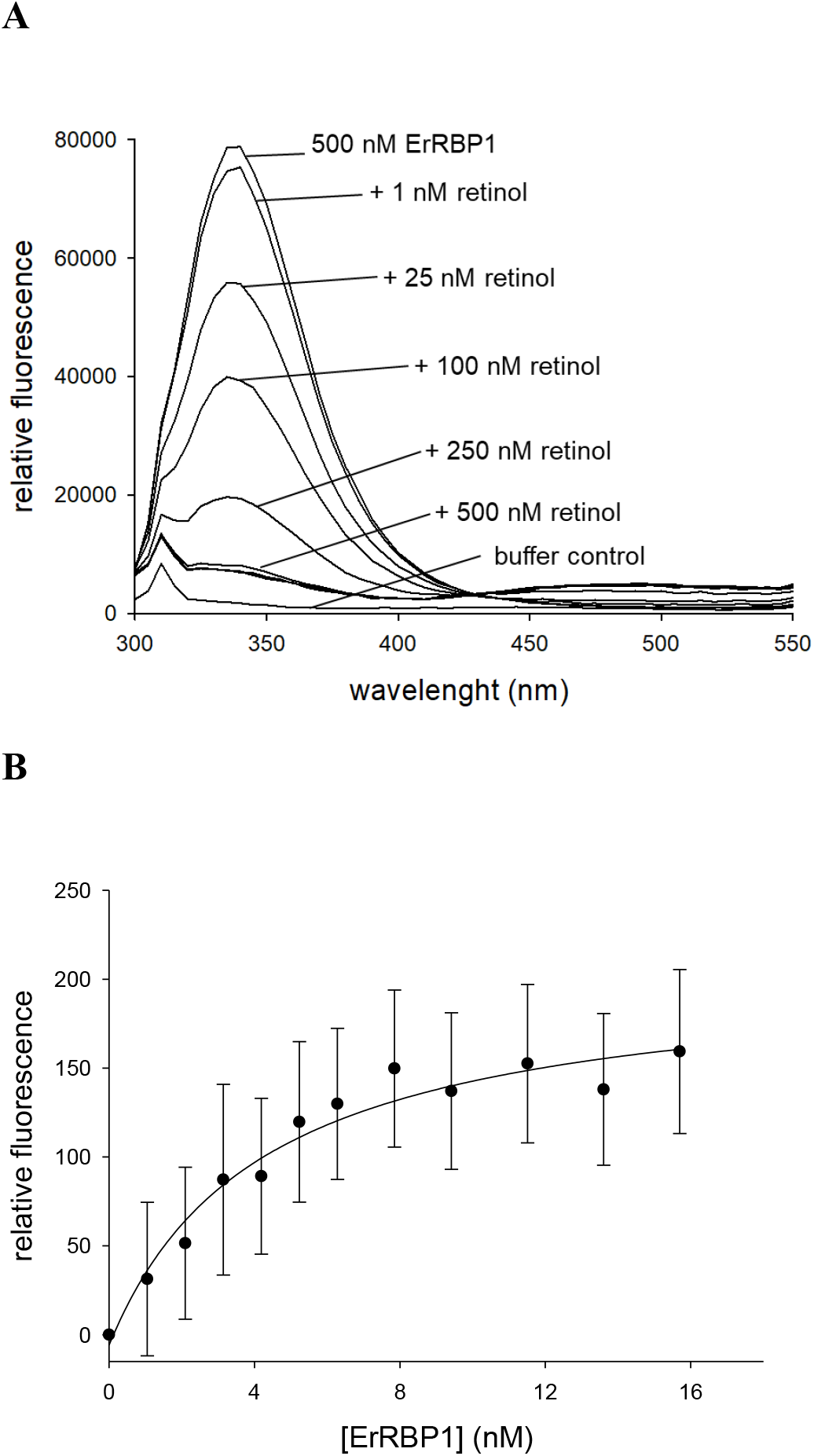
Fluorescence titration of 500 nM ErRBP1 against increasing concentrations of retinol. (**A**) Quenching of the tryptophan fluorescent peak at 340 nm, as well as a visible FRET signal of retinol at 485 nm indicate a binding event. The upper line represents 500 nM ErRBP1, followed by additions of increment steps of retinol as indicated. Additions of 1 µM, 1.5 µM and 2 µM gave nearly identical spectra as seen under the spectrum obtained with 500 nM. (**B**) Binding affinity analysis of the interaction between 7 nM retinol and increasing amounts of ErRBP1. Relative fluorescence emission is measured at 485 nm, excitation wavelength was at 340 nm. The data was fitted by a hyperbolic function yielding an EC_50_ of 4.2 nM.

After we confirmed the expected fluorescence behavior, we aimed at measuring the binding affinity between retinol and ErRBP1 using fluorometric titrations. For this purpose, samples of ErRBP1 and retinol were excited at 340 nm, taking advantage of the substantial increase in retinol fluorescence upon binding to ErRBP1^42^. Emission was monitored at 485 nm over a period of 4 minutes, which greatly increased sensitivity. Binding affinity analyses of the interaction between retinol and ErRBP1 proved challenging, as the data points produced rather segmented and linear curves with a kink at the point of saturation (Figure 2B, can also be noted Figure S2 and S3 in Supplement). Therefore, attempts to fit the curves with exponential binding fits were unsuccessful. However, according to a recent tutorial paper^44^ we reasoned that the concentration of the constant component in our titration experiments (ErRBP1 in Figure 2A, retinol in Figure 2B) is above the actual K_D_. We adjusted the experimental conditions accordingly^44^ by using retinol as the constant binding partner (at 27 nM, 15 nM and 7 nM) and titrated with increasing amounts of ErRBP1 (Figure 2B)^44^. Altogether, the titrations indicated an EC_50_ between retinol and ErRBP1 below 5 nM. The data set shown in Figure 2B was obtained with 7 nM ErRBP1, and a hyperbola curve fitting yielded an EC_50_ of 4.2 nM.

### Protein-protein interaction analysis using SPR

Having established that purified recombinant ErRBP1 is functional, we used the protein for interaction analyses with ErCry4a. Interaction studies with SPR spectroscopy have already been performed using purified ErCry4a when it was shown to interact with the cone-specific G-protein transducin alpha subunit (employed as a Gtα/Giα chimera^31^). The ErCry4a used for this study was sourced from the same lab as the previous study. A detailed protocol of the preparation is available^14^. An example set of new sensorgrams reproducing the interaction between ErCry4a and ErGtα/Giα is shown for internal reference and as a positive control in Figure 3A. Following a similar immobilization protocol and experimental design, we did not observe any specific binding between ErRBP1 and ErCry4a. Although some SPR sensorgrams displayed the typical shapes of association and dissociation events in some recordings (Figure 3 B and C), they failed the critical tests of fulfilling the criteria for a specific binding event. The binding curves did not show a concentration dependency and the maximal amplitudes of the sensorgrams did not correlate with the injected concentration of the analyte (ErCry4a in Figures 3B and C). We also extended the injection time during the association phase enabling us to detect any putative slow association rates that otherwise might have evaded detection. However, the SPR responses showed very low amplitudes that did not vary with the concentration of ErCry4a (Figure 3B). The SPR responses were clearly different from the control recordings in Figure 3A which resulted in a K_D_ of 19.8 nM, which reproduced the high-affinity interaction of ErCry4a with Gtα/Giα reported previously^31^. A direct comparison of ErCry4a binding to either Gtα/Giα or ErRBP1 is shown in Figure 3C, where the sensorgrams are adjusted to the same scale.

**Figure 3.**
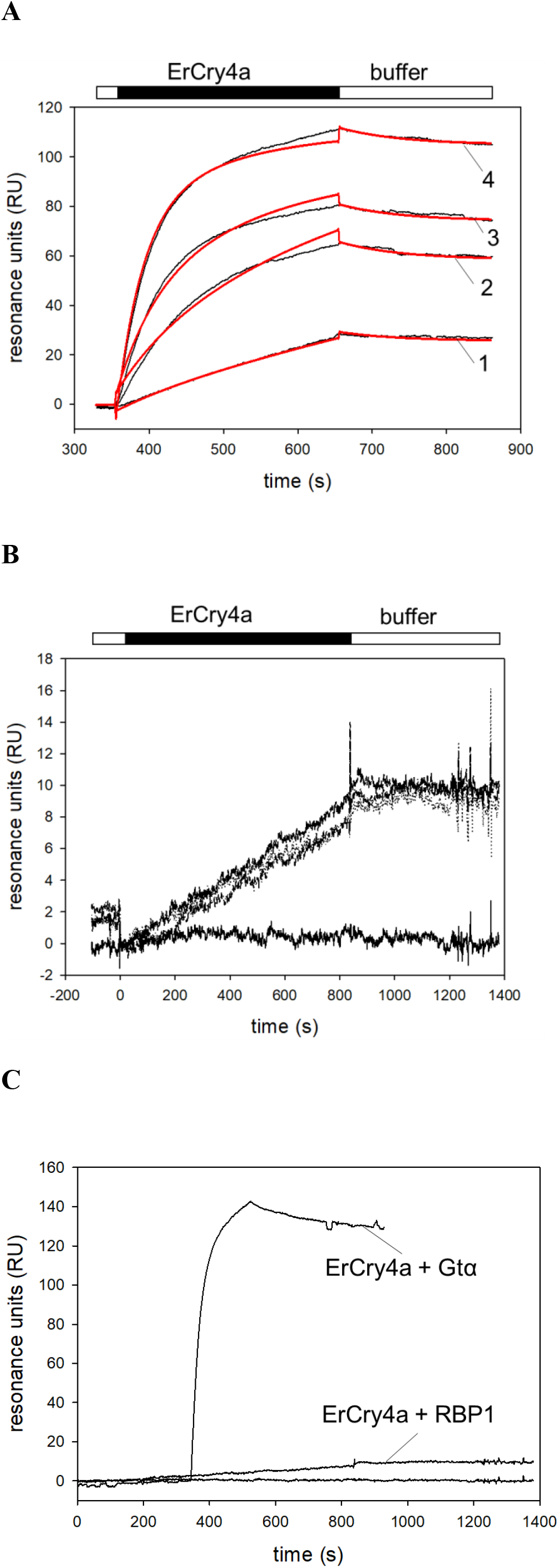
Surface plasmon resonance analysis of ErCry4a protein-protein interactions. (**A**) Binding of *Er*Cry4a (black bar) to non-myristoylated G_t_α/Giα after flushing 5 nM (1), 20 nM (2), 50 nM (3) and 100 nM (4) *Er*Cry4a over immobilized G_t_α/Giα. Global curve fitting (two-state-reaction model, red lines) yielded the following constants: k_a1_ = 2.21×10^5^ M^-1^ s^-1^; k_d1_ = 0.00438 s^-1^ yielding a K_D_ = 19.8 nM. (**B**) Injection of ErCry4a over immobilized ErRBP1. Concentrations of 100 nM, 250 nM and 500 nM ErCry4a all resulted in a nearly identical change in resonance units (upper curves). The lower flat line was obtained by buffer injection. (**C**) Injection of 100 nM ErCry4a over immobilized G_t_α/Giα or ErRBP1 as indicated.

Furthermore, we also tested the interaction between immobilized ErRBP1 and ErCry4a in the presence of retinol. We injected 10 µM retinol in each cycle before the application of ErCry4a. Negative binding curves yielding maximal amplitudes at -20 RU resulted (Figure S4) and showed neither the shape nor the amplitudes seen for the known specific interactions of ErCry4a with Gtα/Giα. In conclusion, we could not detect any strong binding of ErCry4a to immobilized ErRBP1 in the presence of retinol.

### Cellular FRET analysis of ErRBP1 with ErCry4a

However, since RBP1 was identified in a yeast-two hybrid screen, a reasonable question is whether an interaction could occur in a cellular system providing the cytoplasmic milieu of a living cell. We therefore performed a FRET analysis to test for a possible interaction of ErCry4a with ErRBP1 in a neuroretinal quail cell line. We transiently transfected this cell line with ErCry4a labeled with the donor fluorophore mTurquoise and with ErRBP1 labeled with the acceptor fluorophore EYFP to perform acceptor photobleaching FRET. We used both N-terminally labeled (N-protein) and C-terminally (protein-C) labeled proteins and tested all the four different combinations. As a positive control, we again used ErGtα, which we had shown previously to interact with ErCry4a using FRET^31^. This positive control showed that ErGtα and ErCry4a interact in a cellular environment because we observed a higher donor emission after photobleaching of the acceptor, caused by fewer acceptor fluorophores being available for energy transfer than before the photobleaching. The calculated mean FRET efficiency between mTurquoise-ErCry4a (N-ErCry4a) and Gtα-EYFP (Gtα-C) was 24.5 % ± 7.8 % (s.d.). In contrast, FRET efficiencies between N-ErCry4a and N-RBP1 was 1.6 % ± 3.0 % and 1.9 % ± 3.4 % when RBP1-C and N-ErCry4a were used. These FRET efficiencies were in the same range as the negative control N-ErCry4a + EYFP (1.7 % ± 3.2 %). Furthermore, the FRET efficiencies of ErCry4a-C with N-RBP1 (4.9 % ± 4.8 %) and with RBP1-C (4 % ± 3.8 %) were also not significantly different from the negative control ErCry4a-C + EYFP (3.2 % ± 5.2 %) (Figure 4A and B and Table 1). While the absence of FRET cannot rule out an interaction *in vivo*, as the fluorophores could be too far apart or in the wrong orientation to each other, our acceptor photobleaching results further corroborate the absence of an interaction between ErRBP1 and ErCry4a that was also the result of the SPR experiments.

**Figure 4.**
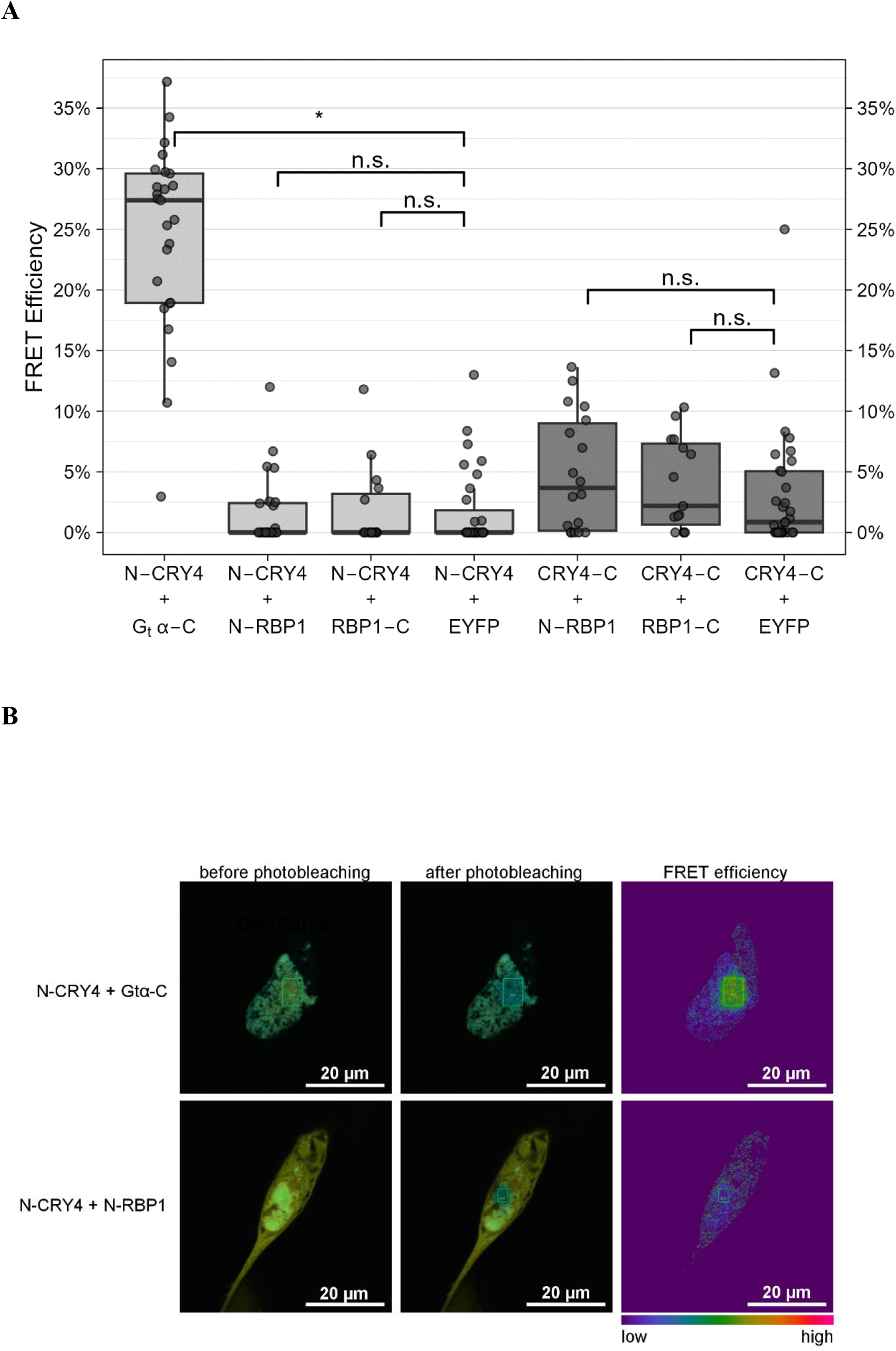
FRET analysis of a possible interaction between ErCry4a and ErRBP1 using the acceptor photobleaching technique. (**A**) FRET efficiencies as determined by the LAS AF software between N- and C-terminally tagged ErCry4a (N-ErCry4a and ErCry4a-C) with mTurquoise as the donor fluorophore and N- and C-terminally tagged ErRBP1 (N-RBP1 and RBP1-C) with EYFP as the acceptor fluorophore. Boxplot graphs were generated using R. The boxes represent the IQR (interquartile range) ranging from Q1 (the first quartile) to Q3 (the third quartile) of the data distributions. Medians are indicated by lines across the boxes. The whiskers extend between the most extreme data points, excluding outliers, which are defined as being outside 1.5 x IQR above the upper quartile and below the lower quartile. Each data point represents the measurement of one cell. Each condition was tested on a minimum of 15 cells in at least three separate experiments. (**B**) Representative confocal laser scanning microscopy images of QNR/K2 cells expressing N-Cry4a + G_t_α-C, used as a positive control, and N-ErCry4a + N-RBP1 before and after photobleaching (overlay of cyan-donor and yellow– acceptor). The green rectangular indicates the bleached area.

**Table 1:** Statistics of FRET monitoring. ANOVA results of the binomial GLM for differences between the interactions of ErCry4a with RBP1 or Gtα and the respective negative controls with EYFP alone. The respective sample sizes (N), resulting chi-squared statistics (χ2), degrees of freedom (Df), and *p*-values as well as Bonferroni-corrected *p*-values are listed for each comparison. Significant differences are indicated with a * (*p ≤ 0*.*05*).

| | N | | N | $\chi^2$ | Df | p-value | corrected p-value | |
| --- | --- | --- | --- | --- | --- | --- | --- | --- |
| **N-CRY4 + Gtα-C** | 25 | N-CRY4 +<br>EYFP | 31 | 7.6177 | 1 | **0.0058** | **0.0173** | * |
| **N-CRY4 + N-RBP1** | 24 |  |  | 0.0004 | 1 | 0.9842 | 1 | n.s. |
| **N-CRY4 + RBP1-C** | 15 |  |  | 0.0025 | 1 | 0.9597 | 1 | n.s. |
| **CRY4-C + N-RBP1** | 18 | CRY4-C +<br>EYFP | 31 | 0.0862 | 1 | 0.7690 | 0.7690 | n.s. |
| **CRY4-C + RBP1-C** | 15 |  |  | 0.0170 | 1 | 0.8962 | 0.8962 | n.s. |

### Discussion

The light-dependent magnetic compass in migratory birds is located in the birds’ retina^5,6,12,45^ and is most likely mediated by Cry proteins^2,3,12,14,27^. Since ErCry4a from the European robin fulfills key biochemical, photochemical, and evolutionary criteria for acting as a magnetically sensitive receptor^14,15,17,24^, the next steps in an ErCry4a triggered signaling pathway await further exploration. Following a lead suggested by Wu et al. (2020)^30^, we investigated whether ErRBP1 directly interacts with ErCry4a and employed different experimental setups. First, we successfully obtained purified ErRBP1 using heterologous expression in *E. coli* and tested its biological function by measuring the binding of retinol. Spectroscopic and fluorescent properties of ErRBP1 (Figure 1 and 2) match previous results obtained with human, rat and chicken cellular retinol binding proteins^41-44, 46-47^. The high affinity binding of retinol to ErRBP1 that we observed, with a K_D_ < 5 nM, is similar to many previously suggested values, which often ranged from ∼100 pM to 16 nM for rat CRBP and human CRBP. We suspect that the differences in the obtained K_D_ values might indicate a species-specific difference or be due to the use of different protocols to evaluate binding affinities as stressed in a recently published tutorial^44^.

Having purified biologically active ErRBP1 in our hands, we could investigate the interaction with ErCry4a on the protein level. In contrast to the yeast-two-hybrid screening approach^30^, we found no experimental evidence for an interaction between ErRBP1 and ErCry4a. We based our observation on two different experimental designs: biosensor technology and cellular FRET measurements in living cells (Figures 3 and 4). These clear negative results were accompanied by positive control experiments and observations (Figures 3A and C, Figure 4), which replicated the results of our previous study confirming that ErCry4a interacts with the cone specific Gα-subunit protein^31^.

However, the presence of retinol might be required for the interaction of ErRBP1 with ErCry4a as it might stabilize the complex formation. Quite surprisingly, our kinetic SPR analysis of ErCry4a flushing over a retinol-saturated ErRBP1 sensor chip surface produced negative response signals (Figure S4) instead of sensorgrams with positive amplitudes that are typical for binding events (Figure 3A and C). Negative SPR signals are rare, but have been observed occasionally, indicating for example, a change in the layer thickness of the dextran sensor surface^48-53^. A change in the hydrodynamic vicinity or water shell of immobilized proteins or conformational transitions can cause such an effect. The presence of retinol might thus change the interaction of ErCry4a with the dextran layer. The five to ten-fold higher concentrations of ErCry4a needed to trigger the response signals in Figure S4 in comparison to Figure 3A (interaction with Gtα/Giα) might point to a very low affinity binding of ErCry4a and ErRBP1. This interpretation might explain why the results related to the Gα-subunit obtained with SPR and FRET approaches matched perfectly the yeast-two-hybrid screening^30^, but did not match in the case of ErRBP1.

Another potential obstacle to studying any binding by SPR might come from the immobilization of ErRBP1 on the SPR sensor chip surface. The immobilization could potentially restrict the accessibility of an interaction domain. We cannot exclude this possibility. One way to avoid such a scenario would be to reverse the immobilization geometry (ErCry4a on the surface), but this procedure will impact on the functionality of ErCry4a and thus the comparability with the results we observed and reported previously^31^. However, any concerns about any problematic SPR immobilization of the target protein do not apply to the FRET measurements, where proteins are freely diffusing in cells. Under these conditions, ErCry4a also bound to the Gα-subunit, but not to ErRBP1 (Figure 4).

Another possibility, which we cannot completely exclude, is that additional proteins might be required for an interaction between ErRBP1 and ErCry4a to take place, and that homologs in yeast were able to fulfill these roles. This seems unlikely, however, as our FRET measurements were performed in a neuroretina cell line from quail, which we think provides the closest possible environment to that where ErCry4a is expected to act as a magnetoreceptor in the European robin. It is difficult for us to believe that yeast was able to provide some hypothetical critical protein partner(s) necessary for this interaction, but our neuroretina cell line from quail was not. However, it should be mentioned that the quail line itself might be limited in its ability to replicate native retinal cells, as it was isolated from a cancerous sarcoma^54^, which likely has a partly altered protein expression profile compared to native retinal cells.

In conclusion, the involvement or participation of a G-protein mediated signaling pathway in magnetoreception seems likely^31,55^. In contrast, a hypothesis involving ErRBP1 is not supported. This is in line with the fact that it is not straightforward to integrate ErRBP1, a protein active in retinoid transport, in a concept of magnetoreceptor signaling. However, ErRBP1 could have a more indirect role in magnetosensory processes. For example, ErRBP1 might have a light-protective role when bound to retinol or operate as an intracellular shuttle for retinol. During these processes, we cannot exclude that low affinity interactions with ErCry4a occur.

## Materials and methods

### Bioinformatics Analysis

Sequences for the homologous RBPs were accessed from the NCBI database (NCBI) using the accession numbers 5H8T_A, P50120, 1GGL_A, Q96R05, P02753, NP_113679, and NP_001264345_XP_422635 for human CRBP1, 2, 3, 4, human RBP 4 and 5, and chicken RBP1, respectively. Additional homologs were identified using the Blast protein suite^56^. Multiple sequence alignments of RBP proteins were performed using Clustal Omega^57^, and estimation of physical protein properties was performed using ProtParam^58^.

### Cloning of ErRBP1

The coding sequence for cellular retinol binding protein 1 from *Erithacus rubecula*^59^ was amplified from cDNA and cloned into a HindIII digested pET-11 vector containing a C-terminal 6x histidine tag using the Gibson assembly. The following primers were used for amplification (5’– GAATTCGAGCTCCGTCGACAAGCTTAGGAGGACAGCTATGCCTGCAGACTTCAAT GG–3’, 5’–GTGGTGCTCGAGTGCGGCCGCAgtCTGCACCTTCTTAAAGACTTGC–3’), and clones were confirmed by Sanger sequencing (Eurofins Scientific).

### Expression and purification of ErRBP1

Chemically competent BL21 cells were transformed using the plasmid and cultivated at 37°C using LB agar plates and liquid media containing kanamycin at 30 µg/mL. Flasks of 700 mL were inoculated and incubated at 37°C with 180 rpm shaking until an OD_600_ of 0.6 was reached, upon which they were induced with 100 µM IPTG. After an additional 4 hours under the same conditions, the cultures were harvested by centrifugation and frozen at -20°C.

Cell pellets from 700 mL culture were resuspended in 15 mL Ni-NTA binding buffer (20 mM Tris pH 8.0, 150 mM NaCl, 10 mM Imidazole, 10 mM ß-mercaptoethanol) supplemented with Lysozyme (0.05 mg/mL), DNAase (0.005 mg/mL), and a protease inhibitor cocktail (Roche - cOmplete™, Mini, EDTA-free Protease Inhibitor Cocktail). Suspensions were then lysed by sonication using a Bandelin Sonopuls Ultrasonic homogenizer at 40% power. Lysates were cleared by centrifugation at approximately 25,000g at 4°C for 60 minutes, and the supernatant applied to a pre-equilibrated 25 mL gravity flow column containing 2 mL of HisPur™ Ni-NTA Resin (Thermo Scientific). Columns were washed with 175 mL of Ni-NTA wash buffer (20 mM Tris pH 8.0, 150 mM NaCl, 20 mM Imidazole, 10 mM ß-mercaptoethanol), and the protein eluted using two 5 mL applications of Ni-NTA elution buffer (20 mM Tris pH 8.0, 150 mM NaCl, 300 mM imidazole, 10 mM ß-mercaptoethanol).

The Ni-NTA elution was prepared for ion exchange chromatography (IEC) by diluting 1:2 in IEC Buffer A (20 mM Tris pH 7.0, 10 mM ß-mercaptoethanol), and was subsequently applied to a HiTrap™ 5 ml Q HP column pre-equilibrated with IEC Buffer A. The column was washed with IEC Buffer A until a stable UV baseline was reached, and the protein was eluted by washing with 95% IEC Buffer A and 5% IEC Buffer B (20 mM Tris pH 7.0, 1 M NaCl, 10 mM ß-mercaptoethanol). The collected protein was either directly used, stored at 4°C, or dialyzed twice against Working Buffer (10 mM HEPES pH 7.4, 150 mM NaCl, 10 mM MgCl_2_, 3.4 mM EDTA) to reach a dilution factor of approximately 100,000, and finally concentrated using a Macrosep Advance 10k MWCO centrifugal filter. Aliquots of protein directly after IEC or dialysis and concentration were then shock frozen in liquid nitrogen and stored at -80° C.

Purity of protein samples was assessed using SDS PAGE using a gel percentage of 15%, followed by Coomassie staining. The molecular weight of the ErRBP1 construct, including the C-terminal 6x histidine tag, was calculated to be 17,051 Dalton based on the amino acid sequence.

### Retinol stock solution

All-*trans*-retinol (Sigma) was dissolved in thoroughly degassed 100% ethanol to a concentration of approximately 10 mM, and small aliquots were stored at -20° C in PCR tubes with minimal headspace. At all stages of work, all-*trans*-retinol (simply termed retinol in the present work) was protected from light by using light-blocking containers and working under red-light conditions when possible.

### ErCry4a Protein Samples

Samples of purified ErCry4a were obtained fresh following the protocol as described^14^, with certain modifications for enhanced efficiency. The LB media composition consisted of 10 g/L yeast extract, and the expression duration was prolonged to 44 hours, deviating from the original 22-hour protocol. Protein purification was performed under dim red light. Immobilized metal affinity chromatography (IMAC) was performed following the procedure detailed by Xu et al.^14^, with a modification in the wash buffer increasing the imidazole concentration from 20 mM to 50 mM. The first purification step was complemented by anion exchange chromatography.

### UV-visible Spectroscopy

Both purified ErRBP1 and retinol were quantified by UV-vis absorption using either a Specord 205 (Analytik Jena) or a Cary 60 (Agilent) double-beam UV-vis spectrophotometer. All final measurements used steps of 1 nm with an integration time of 0.1s between 225-450 nm and were performed in either a 1400 or 3500 µL Quartz SUPRASIL fluorescence cuvette with a PTFE stopper (Hellma). For retinol quantification, mock titrations of the stock solution were performed into 100% ethanol and the concentration determined using a molar extinction coefficient^60^ of 52,800 M^-1^cm^-1^. For ErRBP1, the molar extinction coefficient was estimated to be 26,470 M^-1^cm^-1^ based on amino acid sequence^58^, and was later extrapolated based on biological activity assays.

Four replicate titrations began with ErRBP1 at concentrations between 3.5 and 12 µM dissolved in a 5x dilution of working buffer (2 mM HEPES pH 7.4, 30 mM NaCl, 2 mM MgCl_2_, and 680 µM EDTA). After measuring the protein and buffer baselines, consecutive additions of the retinol stock solution were performed with measurements being made each step. Each titration step was between 1-3 µL in volume, and the total volume titrated remained below 1.5% of the total sample volume. After each addition, the sample was gently but thoroughly mixed by pipetting and measured immediately. Data was fitted linearly, excluding the points closest to saturation, and the intersection of the trendlines was used as the point of saturation.

### Fluorescence Spectroscopy

Retinol absorbs within the wavelengths of intrinsic tryptophan fluorescence at ∼340 nm and subsequently fluoresces at ∼485 nm^42,43^. This pair of excitation and emission wavelengths would allow Förster resonance energy transfer (FRET) measurements suitable for interaction studies. Measurements were recorded on a spectrofluorimeter from Photon Technology International. We confirmed the occurrence of FRET using excitation at 280 nm and recording the emission from 300-550 nm. Fluorometric titrations were performed similarly as described above for UV-vis measurements. Starting with the baseline and measurements of the purified protein alone (∼2.4 µM), retinol was added, and the fluorescence spectrum monitored for changes.

Based on previous observations with human RBP^42^, we used the substantial increase in retinol fluorescence upon binding ErRBP1 to estimate the affinity of the interaction. Several titrations were performed using direct excitation at 340 nm, as this increases the magnitude of the fluorescence signal by avoiding energy loss due to FRET. Samples were measured for a period of 3-5 minutes at 485 nm. These titrations were performed with a fixed retinol concentration followed by stepwise addition of ErRBP1, making the point of saturation clearly recognizable. Replicates were performed with decreasing retinol concentrations to approximately 7 nM retinol. All fluorescence measurements were done in diluted (1:5) IEC buffer at room temperature, with gentle pipetting to mix after each addition, and between 0-2 minutes of incubation time before measuring.

### Surface plasmon resonance (SPR)

SPR protein-protein interaction analysis was conducted on a Biacore 3000 from GE Healthcare, now Cytiva, following the general principles as described previously^31,61^. Experiments were carried out using CM5 sensor chips (GE Healthcare) and the carbodiimide/N-hydroxy-succinimide immobilization technique (Biacore Immobilization Kit, Cytiva, Uppsala, Sweden) for amine coupling of free protein NH_2_ groups, followed by deactivation with 1 M Ethanolamine at pH 8.0. Initial screening of conditions included varying the immobilization geometry (immobilization of ErCry4a with ErRBP1 in mobile phase and vice versa) and density (1600-4200 RU of immobilized protein). An in-house made Ulp1 chain A was used to coat control surfaces with a similar density (within ±15%) to that of the test surface. Final experiments were performed by injecting varying concentrations of purified ErCry4a over immobilized ErRBP1 with flow rates between 15-20 µL/min, contact times between 4-14 minutes, and regeneration with a 60-90 second pulse of a pH 10.25 cocktail containing approximately 15 mM each of ethanolamine, sodium phosphate, piperazine, and glycine, prepared following a previously described protocol^62^. The buffer for all final experiments was 10 mM HEPES/NaOH, pH 7.4, 150 mM NaCl, 10 mM MgCl_2_, 0.005% Tween-20, and 3.4 mM EDTA. We also probed the interaction using ErRBP1 that was saturated with retinol. When ErRBP1 was immobilized, 10 µM retinol was injected in each cycle for 1 min before ErCry4a was flushed over the sensor chip surface. Due to the higher affinity of retinol for ErRBP1 we assumed full saturation of ErRBP1 during the time course of our recordings. All sensorgrams were evaluated using the BIAevaluation software 4.1 (GE Healthcare, Boston, MA, USA).

### FRET analysis in cell culture

*Cloning of FRET constructs –*Both ErCry4a and ErRBP1 were cloned in both an N-terminally tagged version as well as a C-terminally tagged version using the plasmids pmTurquoise-C1 and pmTurquoise-N1 (both a gift from Dorus Gadella (Addgene plasmid # 60558; http://n2t.net/addgene:60558 ; RRID:Addgene_60558 and Addgene plasmid # 60559: http://n2t.net/addgene:60559 ; RRID:Addgene_60559)^63^ for ErCry4a and pEYFP-N1 and pEYFP-C1 (NovoPro Bioscience, Shanghai, China; plasmid #V012015 and #V012016) for ErRBP1. All four plasmids were received as a gift from Prof. Anja Bräuer (Division of Anatomy, University of Oldenburg, Germany). Table S1 in the supplement gives an overview of the cloning details and expressed proteins. In each case, the fluorophores were separated from the cDNA by a linker of 15 to 16 amino acids. Amplifications were done using previously constructed plasmids as templates that contained part of the linker sequence. As a positive FRET control, GNAT2 cDNA coding for G_t_α was also cloned with a C-terminal EYFP tag. Linearized vector and PCR products were recombined using In-Fusion Snap Assembly master mix (Takara Bio, Shiga, Japan).

### Cell Culture and Expression

The QNR/K2 Neuroretina Quail cell line was purchased from ATCC (CRL-2533) and cultured in DMEM + GlutaMAX (Gibco, Waltham, MA, USA) supplemented with 10% fetal bovine serum (Gibco) at 39°C and 5% CO_2_. QNR/K2 cells were (co-)transfected with the FRET constructs using Lipofectamine 2000 (Thermo Fisher Scientific, Waltham, MA, USA) according to manufacturer’s instructions, after which cells were kept in the dark, or handled under dim red light until fixed. Forty-eight hours after transfection, cells were washed twice with PBS (Gibco), fixed in 4% (wt/v) paraformaldehyde for 30 min, and washed again twice with PBS and kept in PBS for imaging. Transfection, fixation and imaging were all performed in chambered coverslips (ibidi µ-Slide 8 Well high with ibiTreat and a #1.5 polymer coverslip bottom (Ibidi, Martinsried, Germany).

*Acceptor photobleaching FRET*—Imaging of transiently transfected QNR/K2 cells was performed with an inverted Leica TCS SP5 II confocal microscope. A 40 x oil objective with 1.3 numerical aperture was used. The donor mTurquoise was excited using an argon laser set to 20 % at 458 nm, and the acceptor EYFP was excited with an argon laser at 514 nm. Photomultiplier tubes were used as detectors. The detection range was set to 465-500 nm for mTurquoise emission and 525-600 nm for EYFP emission using AOTF (acousto-optical tunable filter). To determine the FRET efficiency, the acceptor photobleaching protocol was used. Acceptor photobleaching was performed with the FRET AB wizard in the LAS AF software on selected regions of interest (ROIs), where both fluorophores showed expression. Acceptor bleaching was achieved by using 70 % intensity of the argon laser at 514 nm and two bleaching frames. Prebleach and postbleach images were acquired. FRET for the ROIs was observed by an increase in donor (mTurquoise) fluorescence intensity following the acceptor (EYFP) photobleaching. FRET efficiency was measured in percent and calculated automatically by Leica LAS AF software as (Dpost - Dpre)/Dpost, where Dpost (Dpre) is the fluorescence intensity of the donor after (before) photobleaching. For each condition, at least three individual experiments from freshly transfected cells were performed.

*Statistics* — Statistics on FRET data was done as previously^31^ using a binomial generalized linear model (GLM) followed by an analysis of variance, type III from the ‘car’ package in reference^64^ of the resulting models to compare interactions and negative controls. The data were analyzed with a custom-written R-script^65^.

## Acknowledgements

KWK. and HM gratefully acknowledge funding from the DFG research training group grant GRK 1885/2, from the DFG SFB 1372 “Magnetoreception and navigation in vertebrates” (No. 395940726) and from an AFOSR grant (No. FA9550-14-1-0095). HM acknowledges funding from the European Research Council (under the European Union’s Horizon 2020 research and innovation programme, grant agreement no. 810002 (Synergy Grant: “QuantumBirds”). The authors acknowledge the Fluorescence Microscopy Service Unit, Carl von Ossietzky University of Oldenburg, for the use of the imaging facilities.

## Author contributions

CY, RB and KWK conceived and supervised the study. CY, RB, KG and JS performed experiments and collected data. BL performed the statistical analysis. All authors analysed data. CY, RB and KWK wrote the draft of the manuscript. All authors corrected the manuscript. All authors reviewed and approved the final version of the manuscript.

## Data availability statement (mandatory)

All data are available in the main text and /or the supplementary materials. Additional data to this paper may be requested from the authors.

## Additional Information (including a Competing Interests Statement)

All authors declare no competing interest.

