## Supplementary material for "Comparison of retinol binding protein 1 with cone specific G-protein as putative effector molecules in cryptochrome signalling"

### to whom correspondence should be addressed:

#### Table S1

*Table S1: Overview of the cloning details of the plasmids used in the FRET measurements*

| vector | restriction sites used | cDNA | Forward primer | Reverse primer | Amino acid linker | Plasmid name | Expressed protein (short name) |
|---|---|---|---|---|---|---|---|
| pmTurquoise-N1 | *Xho*I + *Apa*I | *Er*Cry4 | GGACTCAGATCTCGAGCCACCATGCTGCATCGCACCAT | GACCGGTGGATCCCGGGCCTCGCCGCCGCTTTCC | LELESGGEARDPPVAT | p*Er*Cry4-mTurquoise | CRY4-C |
| pmTurquoise-C1 | *Xho*I + *EcoR*I | *Er*Cry4 | GGACTCAGATCTCGAGGATCCTCGGGATCATCAGG | GTCGACTGCAGAATTCTTATTCTGTTGTTCGGGCCACTT | SGLRSRGSSGSSGAP | pmTurquoise-*Er*Cry4 | N-CRY4 |
| pEYFP-N1 | *Xho*I + *Apa*I | *Er*RBP1 | GGACTCAGATCTCGAGCCACCATGCCTGCAGACTTCAATGGG | GACCGGTGGATCCCGGGCCTCgccgccgctttcc | LELESGGEARDPPVAT | p*Er*RBP1-EYFP | RBP1-C |
| pEYFP-ErCry4 | *Asc*I + *Apa*I | *Er*RBP1 * | | | SGLRSRGSSGSSGAP | pEYFP- *Er*RBP1 | N-RBP1 |
| pEYFP-N1 | *Xho*I + *Apa*I | *Er*GNAT2 | GGACTCAGATCTCGAGCCACCATGGGGAGCGGGGC | GACCGGTGGATCCCGGGCCTCGCCGCCGCTTTCC | LELESGGEARDPPVAT | p*Er*$G_t\alpha$ -EYFP | $G_t\alpha$-C |
| pEYFP-C1 | *Xho*I + *EcoR*I | *Er*Cry4a | GGACTCAGATCTCGAGGATCCTCGGGATCATCAGG | GTCGACTGCAGAATTCTTATTCTGTTGTTCGGGCCACTT | SGLRSRGSSGSSGAP | pEYFP-*Er*Cry4 | not used in this study |
| pKanCMV-mRuby3 | *Bsp*1407I + *Apa*I | *Er*RBP1 | ATGGACGAGCTGTACAAGAGTGGATCCTCGGGATCATCAGGCGCGCCTATGCCTGCAGACTTCAATGGG | GACACTATAGAATAGGGCCCTCACTGCACCTTCTTAAAGACTTGCTTACATG | KSGSSGSSGAP | pKan-mRuby3-*Er*RBP1 | not used in this study |

*directly cut from pKan-mRuby3-*Er*RBP1

**Figure S1**

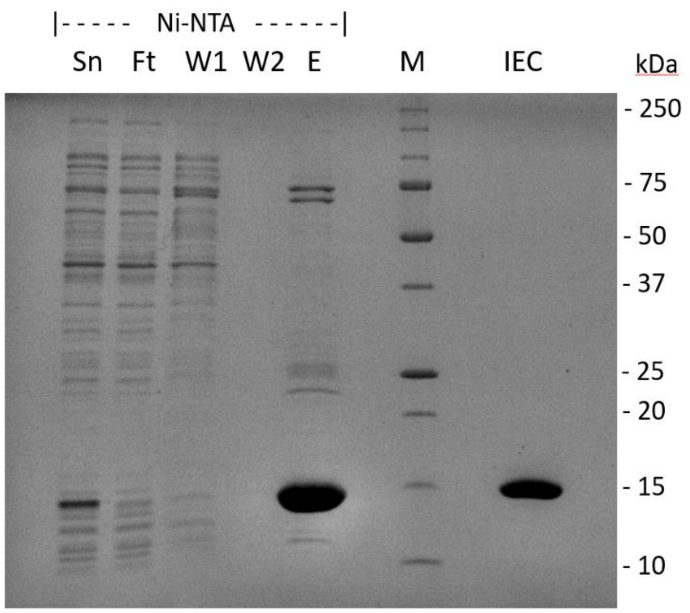


**Figure S1.** Purification of ErRBP1 analyzed by SDS PAGE. Coomassie staining of a 15% polyacrylamide gel showing selected purification steps: M – Marker (corresponding molecular masses are indicated on the right), Sn – Supernatant, Ft – Flow through, W1 – Wash: first 25 mL, W2 – Wash: last 25 mL, E – Elution, IEC – collected IEC fraction. The collected IEC fraction was used for experiments.

#### Titration of ErRBP1 with all-trans-retinol

The characteristic change in the absorbance spectrum of retinol upon binding RBP1 allowed us to extrapolate the amount of active protein in the sample, and to experimentally measure the molar extinction coefficient using a simple UV-vis assay. For this assay, an initial starting concentration of ErRBP1 was quantified at 280 nm (bold black line in Figure S2), and then directly used for a titration using carefully defined amounts of retinol. As retinol is added, the characteristic spectrum of RBP-bound retinol is clearly visible, and the spectra increase in magnitude linearly according to the retinol concentration (green lines in Figure S2). Upon reaching saturation, the character of these additions abruptly changes, as no more RBP is available for retinol binding. The additions post-saturation also increase in magnitude linearly, and often with a slope differing from that of the pre-saturation additions. By plotting single wavelengths within the spectra vs. retinol added, we found linearly increasing values with a sharply defined kink at the point of saturation (Figure S3). The kink was most pronounced for the wavelength of 368 nm, and this wavelength was thus preferred in this study. Finding the intersection of linear fits pre- and post-saturation then led us to point of saturation and the correlating retinol concentration. Using this concentration and the original protein absorbance at 280 nm, we calculated the experimental molar extinction coefficient using the Beer-Lambert law under the assumption of a 1:1 binding stoichiometry, 100% biological activity, and the absence of contaminants absorbing at 280 nm. After four replicates with initial RBP concentrations ranging between 3.5-12 µM, we suggest an experimental molar extinction coefficient for ErRBP1 at 280 nm of 25560 ± 1037 $M^{-1}cm^{-1}$. A breakdown of the approach described above is shown below.

*A = εℓc (Beer-Lambert Law)*

*A* is the absorbance (of ErRBP1 sample at 280 nm pre-titration)

*ε* is the molar attenuation coefficient or absorptivity of the attenuating species

*ℓ* is the optical path length

*c* is the concentration of the attenuating species (concentration of retinol added at point of saturation, identified using absorbance values at 368 nm). Further extrapolating on other wavelengths within the data from Figure S2, we calculated a molar extinction coefficient for retinol bound to ErRBP1 of 50,960 ± 2690 $M^{-1}cm^{-1}$ at 350 nm, and that for free aqueous retinol of ~26,100 ± 1045 $M^{-1}cm^{-1}$ at 325 nm, a mere ~49.5 ± 4% of the original value for 100% ethanol published in Garwin and Saari (2000)[60].

**Figure S2**

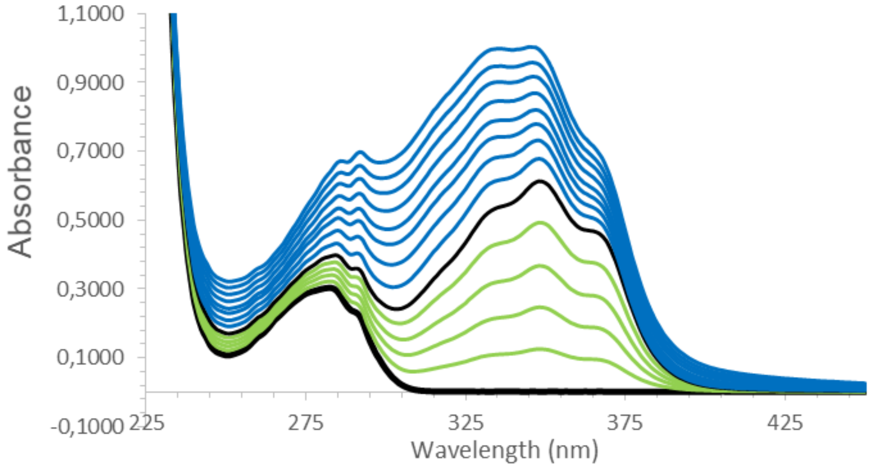


**Figure S2.** UV-vis Absorbance spectrum of purified ErRBP1 and consecutive additions of retinol. The protein concentration was estimated to be 11.3 µM, and retinol concentrations are shown at increasing intervals of 2.32 µM up to a total of 30.3 µM. A thicker line is shown where the titration approximately reaches saturation (11.6 µM retinol).

**Figure S3**

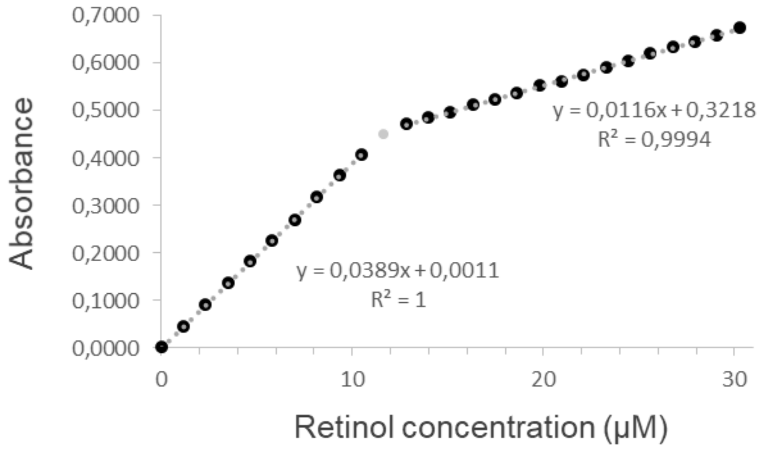


**Figure S3.** Evaluation of the UV-vis absorbance spectra shown in Figure S2. Absorbance at 368 nm is determined and presented as a function of the retinol concentration. Linear fits before and after saturation is reached allow for estimating the amount of active protein.

**Figure S4**

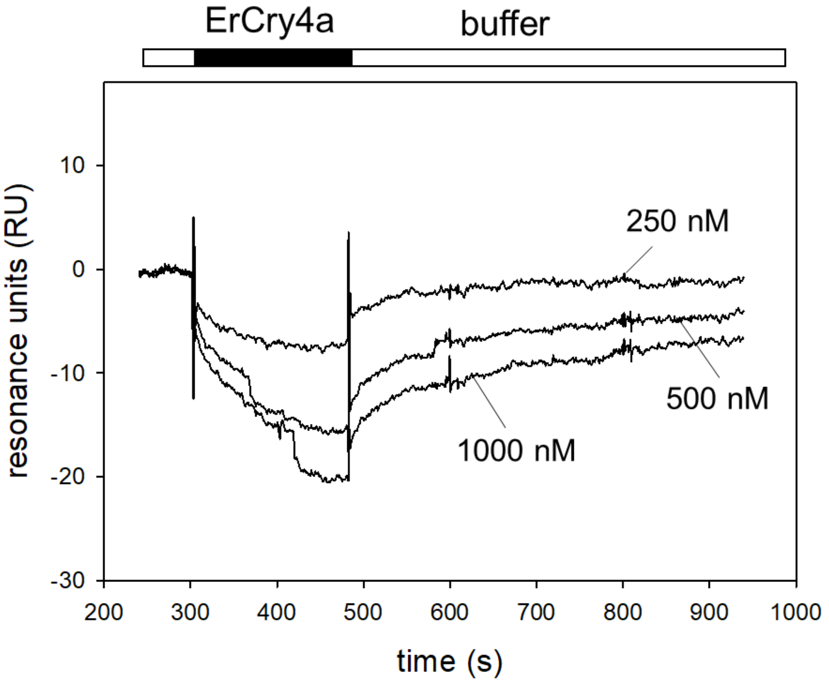


**Figure S4.** Injection of ErCry4a over immobilized ErRBP1 saturated with retinol. Concentration of 250 nM, 500 nM and 1000 nM ErCry4a resulted in negative responses of low amplitudes.
